# Amyloid Explorer: a global atlas of amyloid fibril structures and thermodynamic principles

**DOI:** 10.1101/2025.10.15.682595

**Authors:** Vasilis Kyriazis, Laxmikant Gadhe, Katerina Konstantoulea, Nikolaos Louros, Joost Schymkowitz, Frederic Rousseau

**Author notes:** Correspondence should be addressed to: N.L, J.S, F.R.

## Abstract

Amyloid fibrils underpin both functional and pathological protein assemblies and exhibit extensive structural polymorphism. More than 600 high-resolution fibril structures are now available across over 50 protein sequences, yet comparative analysis has been hindered by fragmented resources and inconsistent annotation. Here we present Amyloid Explorer, an open-access platform that integrates the complete fibril archive into a standardized, quality-controlled, and thermodynamically annotated framework. Each entry includes residue- and segment-level stability maps, structural quality metrics, and interactive visualization tools. Global analysis across the dataset reveals conserved rules of amyloid assembly, including cooperative cores, framework polymorphism through reuse of conserved segments in distinct folds, polymorph-specific energetic anchors, and short frustration zones. By combining scale with accessibility, Amyloid Explorer transforms a static archive into a discovery platform for mechanistic insight, classification, and the design of aggregation modulators.

## Introduction

Amyloid fibrils are highly ordered protein assemblies defined by a cross-β architecture and are central to a wide spectrum of biological and pathological phenomena. They are implicated in neurodegenerative disorders such as Alzheimer’s, Parkinson’s, and systemic amyloidoses^1-3^, but also serve physiological roles in bacterial biofilms, hormone storage, and memory-related processes^4-6^. In recent years, cryoelectron microscopy and solid-state NMR have provided an explosion of atomic-resolution amyloid fibril structures, revealing extensive polymorphism across different proteins and experimental origins, including fibrils assembled in vitro from recombinant proteins and those extracted directly from patient-derived material^7-11^.

More than 600 amyloid fibril structures have now been deposited in public databases, covering over 50 protein sequences. Yet despite their number and quality, these structures remain difficult to compare or interpret. Coordinate files do not readily expose the stabilizing interactions within fibrils, indicate model reliability, or reveal the molecular determinants of polymorphism. Moreover, while structural repositories such as the Protein Data Bank (PDB)^12^, the Electron Microscopy Data Bank (EMDB)^13^, and the Biological Magnetic Resonance Data Bank (BMRB)^14^ provide essential access to raw data, they lack fibril-specific annotations and analytic tools to support mechanistic interpretation^15,16^. As a result, much of the insight embedded in the expanding fibril structure archive remains untapped.

Several recent efforts have begun to chart the growing amyloid fold space by cataloguing individual polymorphs or comparing select fibril families^16-19^. Others have explored local energetics using simulation-driven thermodynamic scans^19-23^ and topological clustering of fibril geometries^24^ or reviewed emerging biophysical principles^25,26^. A number of databases have also emerged, ranging from sequence-centric repositories such as Waltz-dB^27^, AmyPro^28^ and CPAD^29^ to structural platforms such as the Amyloid Atlas^26^, and STAMP-dB^30^. The latter provided the first systematic demonstration that integrating energetic annotation with structural data can illuminate patterns of stability and frustration in amyloid fibrils. Despite these advances, existing resources remain restricted in scope (e.g., to pathological human amyloids), emphasize sequence-level annotations, or lack systematic thermodynamic profiling and cross-fibril comparison. Consequently, the wealth of information embedded in the rapidly growing structural archive remains difficult to access and interpret in a unified way.

Here, we present Amyloid Explorer, a comprehensive resource that systematically integrates all available amyloid fibril structures, currently encompassing more than 600 structures, into a single analytical framework. The platform provides with analytical thermodynamic annotation, structural quality metrics, and comparative visualization tools. This allows for local structural analyses, as well as supports a first extraction of generalizable amyloid architectural rules, enabling global comparisons akin to the early mapping of globular protein folds in the 1980s. Beyond serving as a structural repository, Amyloid Explorer functions as a discovery engine. Comparative analyses across the full dataset reveal generalizable features of amyloid architecture, including (i) the recurrent reuse of aggregation-prone regions (APRs) as stabilizing cores, (ii) polymorph-specific rearrangements of a limited set of stabilizing “anchor” residues, and (iii) the localization of frustrated or destabilizing segments at protofilament interfaces. Together, these findings suggest that amyloid fibrils, despite their apparent diversity, adhere to a constrained thermodynamic framework shaped by recurring energetic and structural principles. This framework provides a foundation for predicting polymorph behavior, understanding strain specificity, and guiding the rational design of modulators or inhibitors.

## Results

### Fibril structure coverage and dataset standardization

Amyloid Explorer compiles structural data from over 600 experimentally determined amyloid fibril assemblies, representing more than 40 distinct proteins (Fig. 1A-C). These include well-studied pathological fibrils such as Aβ, tau, α-synuclein, and TDP-43, as well as more recently characterized functional systems including antimicrobial peptides, hormone-associated amyloids, and synthetic designs^31-35^ (Fig. 1D). The dataset captures broad biochemical and experimental diversity, encompassing fibrils assembled in vitro from recombinant proteins, seeded assemblies, and patient-derived ex vivo material (Fig. 1E-F). The majority of structures were resolved by cryo-electron microscopy (≈ 90%), with the remainder derived from solution and solid-state NMR (Fig. 1G).

**Figure 1.**
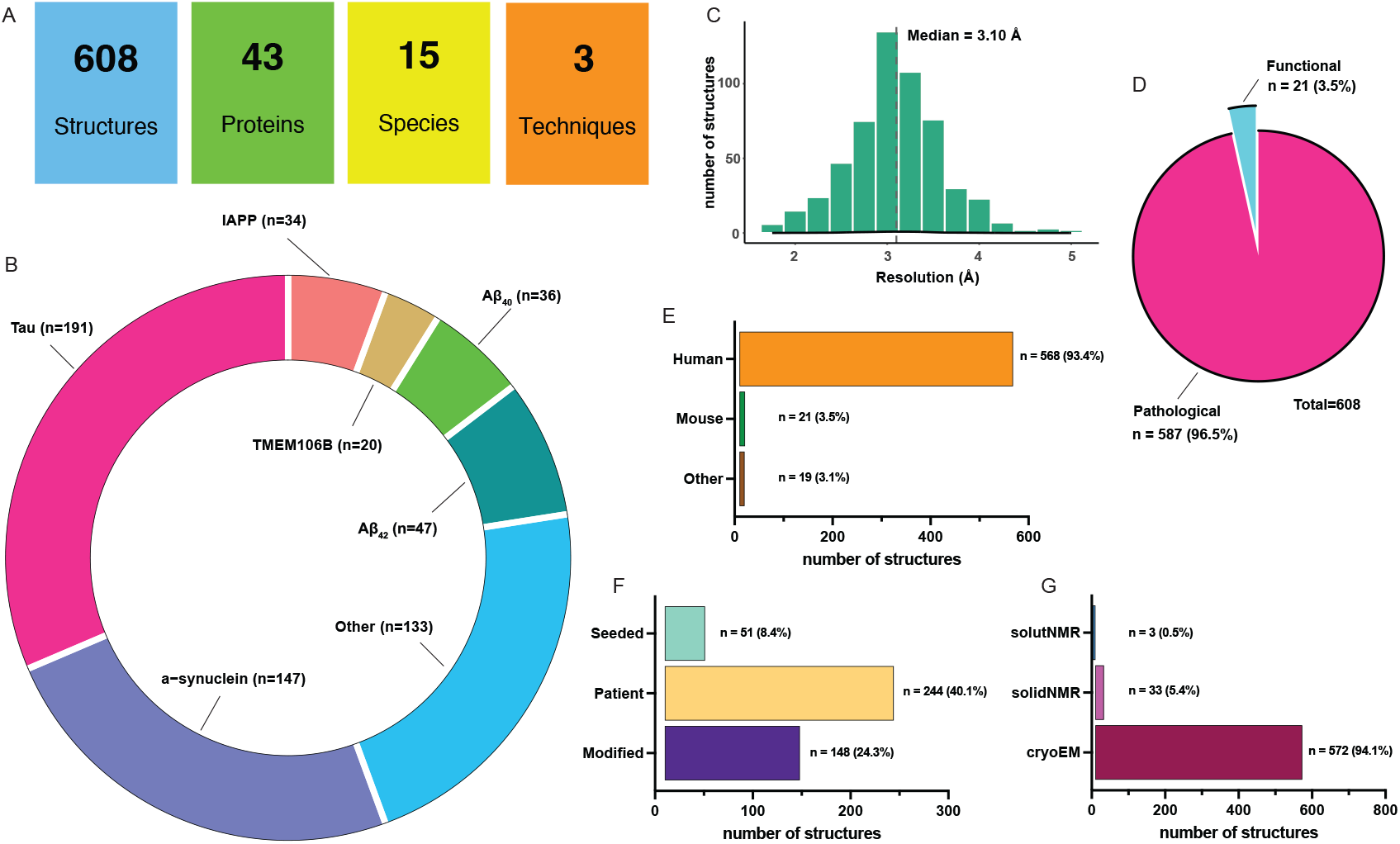
Overview of the Amyloid Explorer structural dataset. **(A)** Summary statistics showing the total number of deposited amyloid structures, unique proteins, represented species, and experimental techniques. **(B)** Donut chart highlighting the dominant proteins for which multiple fibril structures have been solved to date. **(C)** Distribution of structural resolutions across all entries, with the median resolution (3.1 Å) indicated by a red dashed line. **(D)** Pie chart illustrating the relative representation of functional versus pathological amyloid structures. **(E)** Organismal distribution of structures (human, mouse, and other species). **(F)** Bar chart showing the number of structures derived directly from patient samples or seeded with patient material, as well as those containing post-translational modifications (PTMs), insertions/deletions, or disease-associated mutations (Modified). **(G)** Distribution of structures determined by each experimental method (cryo-EM, solution NMR, and solid-state NMR).

To enable direct thermodynamic and structural comparisons, all models were processed to contain ten stacked monomers per protofilament and consistently renumbered. Protofilament chains were renamed and aligned to standardized conventions, and heterotypic components (e.g., detergents or cofactors) were removed when incompatible with energetic calculations. This preprocessing enables residue-level and cooperative window-based ΔG annotation across assemblies and allows comparison of sequence coverage, polymorphic diversity, and energetic architecture within and across proteins.

The resulting dataset reflects a sixfold expansion in fibril coverage relative to our previous implementation of STAMP-dB^30^, and is continuously updated as new structures become available. Importantly, the database includes not only structural coordinates but also metadata annotations covering experimental origin, resolution, mutational status, post-translational modifications, functional assignment (pathological vs. functional), and links to source PDB^12^, EMDB^13^, BMRB^14^ archives and primary publications.

### Interactive data access and visualization

Amyloid Explorer is designed as an open-access, interactive platform that enables users to explore and analyze fibril structures both individually and across the full dataset. The database home page introduces the scope, structure types, and recent updates. From there, users can access a searchable and filterable table of all curated fibril entries (Fig. 2), sortable by protein name, resolution, experimental origin (recombinant, ex vivo, or seeded), polymorph count, and functional assignment. A dynamic sidebar enables rapid subgrouping by key metadata fields such as disease mutation status or post-translational modifications.

**Figure 2.**
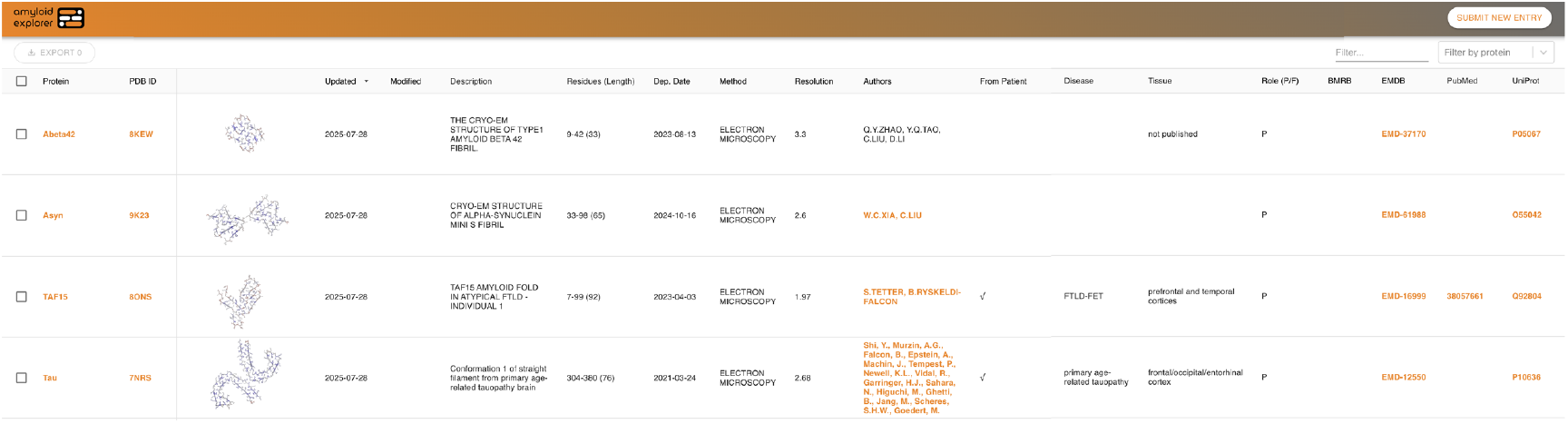
Amyloid Explorer database overview. The database view lists all curated fibril structures in a sortable and filterable table. Each row includes protein name, PDB ID, experimental origin (recombinant, ex vivo, or seeded), method, resolution, mutation and PTM status, tissue source (if applicable), structural links (PDB/EMDB/BMRB), and other metadata. Downloadable cross-section thumbnails offer an immediate visual cue of protofilament architecture. The left-hand protein shortlist and filter bar allow for rapid subsetting. An export button enables instant download of filtered datasets for downstream analysis.

Each entry links to a dedicated protein page that collates all available fibril polymorphs for that sequence. A sequence-level panel shows the extent of each protofilament, with energetic coloring according to per-residue or sliding-window ΔG (Fig. 3A). Disease mutations, deletions, and aggregation-prone regions extracted from literature are annotated directly on the sequence bar. A toggle allows switching between energy modes and highlights differences between polymorphs.

**Figure 3.**
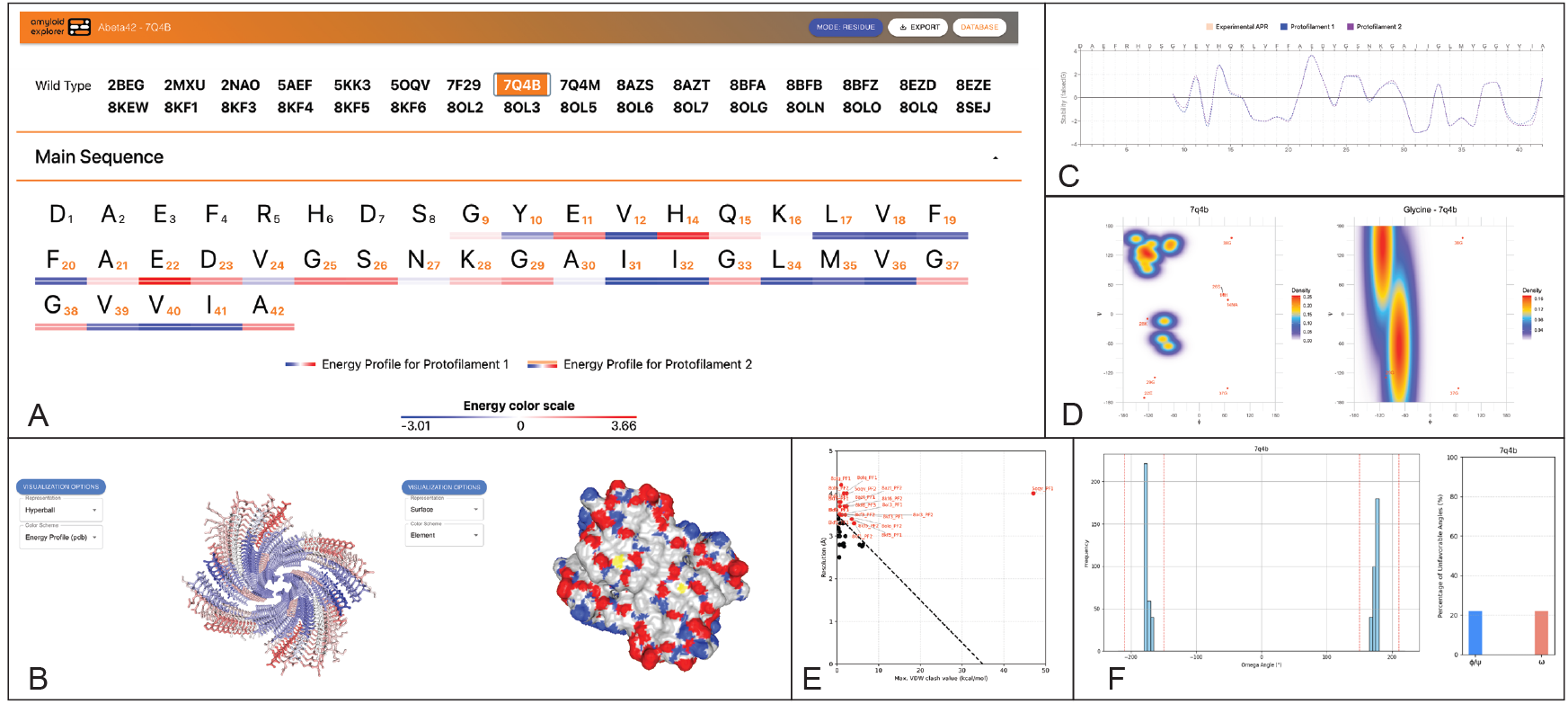
Protein entry page features and analysis panels. **(A)** Header and sequence panel for a selected protein entry, showing PDB/EMDB identifiers grouped by wild-type and variant structures. The energetics toggle (highlighted in red) switches between residue-level and sliding-window ΔG profiles. Annotated sequence bar indicates the fibril core per protofilament and overlays disease mutations, post-translational modifications, deletions, and literature-defined aggregation-prone regions. **(B)** Interactive 3D viewer showing a protofilament structure in cartoon and surface representations. Color schemes can be applied based on energetics, hydrophobicity, secondary structure, chain ID, or residue index. **(C)** Thermodynamic profile plot of ΔG across the sequence for two protofilaments. Experimental APRs are shaded, and the plot synchronizes with the selected energetic mode. **(D)** Ramachandran plot distributions per protofilament, with glycine-specific contours shown separately. **(E)** Clash-versus-resolution scatterplot highlighting fibril structures that exceed the empirical quality threshold. **(F)** Histogram of ϕ/ψ and ω dihedral angles and outlier distributions across protofilaments.

An embedded 3D viewer^36^ allows users to explore individual structures using multiple representations (cartoon, surface, spacefill, ball-and-stick) and coloring schemes based on thermodynamic values, secondary structure, hydrophobicity, or residue index (Fig. 3B). Real-time toggling between energetic modes enables visual comparison of structural stabilization across polymorphs. Below the viewer, an interactive plot displays ΔG profiles per protofilament, with APRs highlighted for reference (Fig. 3C).

Users can export sequence-annotated energetic profiles, structure visualizations, and filtered protein lists in multiple formats. All entries are directly linked to source records in the PDB, EMDB, and BMRB, as well as to primary literature references. The platform requires no login and supports programmatic access through documented API endpoints.

### Structural quality assessment

Each fibril model in Amyloid Explorer undergoes systematic quality control to facilitate reliable downstream analysis and user-driven filtering. Ramachandran plots (Fig. 3D) generated using internal scripts and PyRosetta^37^ are available for each polymorph. Resolution values are extracted from original PDB or EMDB metadata, typically based on Fourier Shell Correlation (FSC) 0.143 thresholds reported in the source publications^38^. Van der Waals (VDW) clashes are quantified using the FoldX force field^39^, and plotted against reported resolution to flag models with disproportionately high steric overlap relative to map quality. To assist structural validation, each protein entry includes a clash-versus-resolution scatterplot (Fig. 3E), highlighting models that exceed an empirical quality boundary derived from prior benchmarks^17^. Additional histograms display the distribution of ω dihedral angles and the fraction of φ/ψ and ω outliers per protofilament, segmented by residue class (glycine). φ/ψ and ω outlier percentages are summarized per model (Fig. 3F).

These quality metrics are embedded directly into the database interface, allowing users to filter entries by resolution thresholds, steric clash rates, or backbone geometry. This enables selective exploration of high-confidence models and minimizes confounding noise in comparative energetic analysis.

### Thermodynamic annotation

Amyloid Explorer provides dual-mode thermodynamic profiling for each fibril structure, offering both residue-level and sliding-window free energy maps. These annotations reveal localized stabilizing and destabilizing elements and enable the identification of cooperative aggregation-prone regions (APRs).

Per-residue ΔG values are computed using FoldX^39^, assessing each residue’s energetic contribution to protofilament stability and cross-β packing. Sliding-window ΔG is calculated using a previously validated cooperative model^17^, which captures multi-residue stabilizing segments and has been shown to align with aggregation-prone cores validated by hydrogen-deuterium exchange, limited proteolysis, and other biophysical data^40-42^.

These two thermodynamic views are available as a toggle in each protein entry and are color-mapped onto both the sequence and the 3D structure (Fig. 3A–C). Sliding-window maps are particularly useful for visualizing cooperative cores, while residue-level data provides finer granularity at polymorphic interfaces or mutation sites. Energetic profiles are also plotted below the structure viewer, with APRs highlighted for reference and plotted separately for each protofilament to visualize symmetry or asymmetry in stabilization.

This dual annotation strategy enables identification of stabilizing hotspots, frustrated regions, and energetic asymmetries within and across fibrils. The integrated presentation of sequence, structure, and energetics allows users to map disease mutations onto energetic landscapes, compare fibril variants, and infer structure–function– stability relationships.

### Global thermodynamic principles of amyloid polymorphism

To connect embedded energetics to composition and physicochemical descriptors, we asked whether residue-level ΔG values, compiled consistently across structures, could reveal global regularities that transcend individual proteins. Aggregating per-residue values across the dataset, we first examined amino-acid-wise mean energetic contributions. The rank order follows classical intuition: hydrophobic residues are, on average, more stabilizing, whereas polar and charged residues tend toward neutral or destabilizing contributions (Fig. 4A). We then systematically correlated these mean per-residue energies with a comprehensive panel of physicochemical scales from AAindex^43^. Volcano analyses and multiple-testing-corrected correlations highlighted a compact set of the strongest associations (Fig. 4B). Hydrophobicity-(GOLD730101) and side-chain property scales (GRAR740102), together with aggregation propensity as captured by Aggrescan^44^, showed robust linear relationships with the embedded energetic estimates (Fig. 4C–E). These associations provide a direct bridge between structure-embedded energetics and established biophysical descriptors, reinforcing the physical interpretability of the ΔG maps.

**Figure 4.**
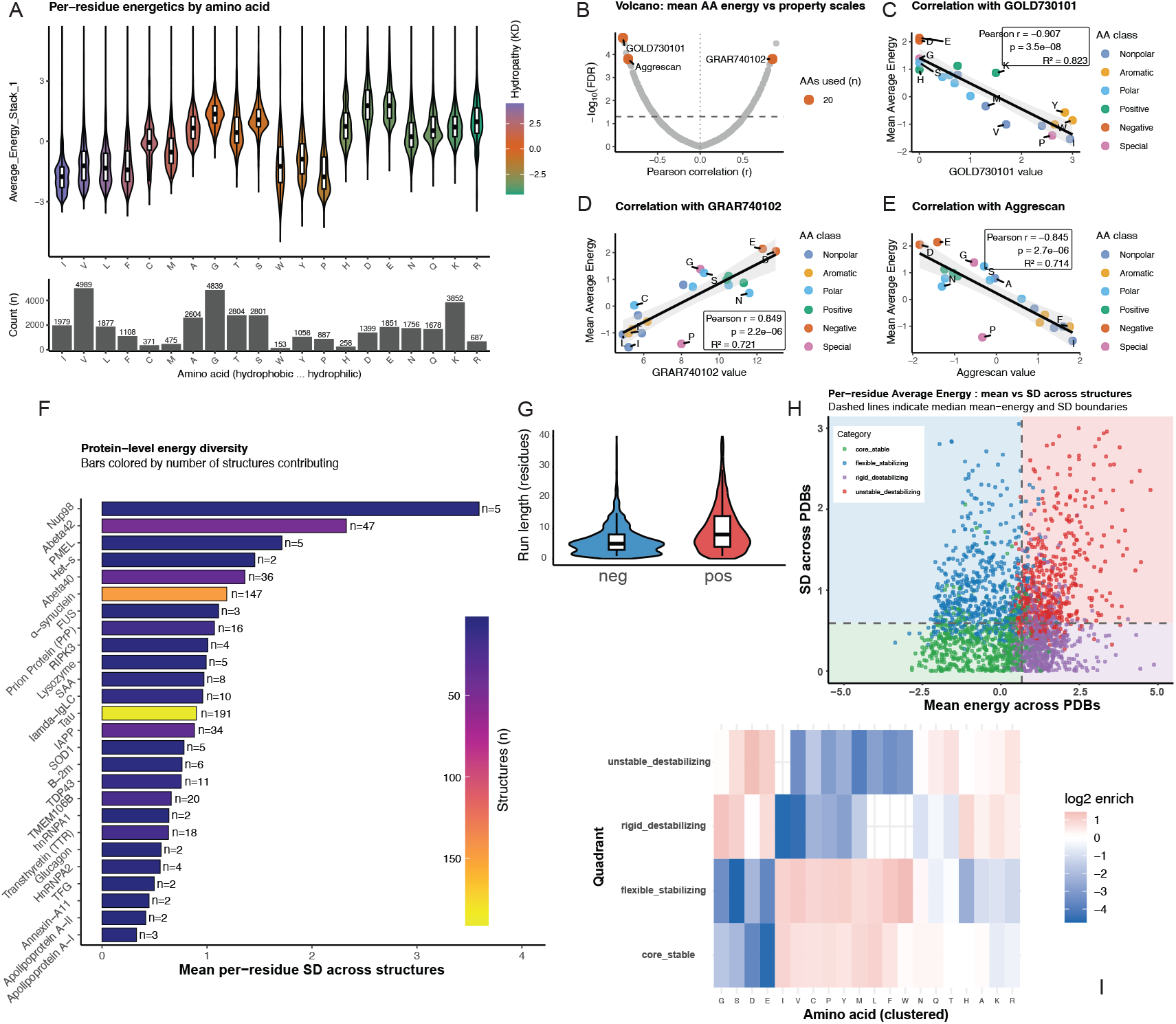
Thermodynamic rules of amyloid structural polymorphism. **(A)** Per–amino-acid energetics. Violin/box plots of per-residue energy values grouped by amino-acid type and ordered by Kyte–Doolittle hydropathy; the hydropathy gradient is shown on the violins. The aligned lower panel reports sample sizes (n) for each amino acid. (B–E) Scale–energy relationships. **(B)** Volcano plot ranking the correlation between mean per-AA energy and hundreds of physicochemical scales (effect size = Pearson r, significance = −log10 FDR). **(C-E)** Representative correlation panels for three selected scales, with least-squares fit, 95% CI shading, and selective residue labels. **(F)** Protein-level energy diversity. Mean per-residue SD across structures (one value per protein), with bars colored by the number of structures contributing to each estimate and the corresponding size (n) annotated at bar ends. **(G)** Stabilizing vs. destabilizing region lengths. Violin/box plots of run lengths for negative-energy (“stabilizing”) versus positive-energy (“destabilizing”) stretches (from Avg_SW_Energy_Stack_1), pooled across structures. **(H)** Global quadrant map. Scatter of perresidue mean energy versus SD across structures, colored by quadrant (core-stable, flexible-stabilizing, rigid-destabilizing, unstable-destabilizing) with global median cutlines shown as dashed guides and faint quadrant backgrounds for interpretability. **(I)** Quadrant-specific composition. Heat map of log2-enrichment of amino acids within each quadrant relative to the global background; amino acids are clustered by their enrichment pro les to emphasize patterns shared across quadrants.

Because amyloid polymorphism is a property of proteins rather than residues, we next summarized heterogeneity at the protein level. We defined an energy-diversity score as the mean standard deviation of per-residue ΔG across all available structures for each protein. This measure varied widely across proteins (Fig. 4F). Visual encodings that reflect the number of structures contributing to each estimate allow one to separate genuine energetic heterogeneity from the appearance of variability introduced by sparse sampling, thereby highlighting proteins whose landscapes have inherent plasticity compared to those with more constrained energetic organization.

Recognizing that stabilization and frustration are often realized as runs rather than isolated sites, we compared the length distributions of contiguous negative-ΔG (stabilizing) and positive-ΔG (destabilizing) segments. The resulting distributions are distinct: Stabilizing segments are short, with mean run lengths around five to six residues, consistent with the typical lengths of compact cooperative cores/APRs, while destabilizing segments are broader and enriched at structural boundaries and putative co-factor interfaces (Fig. 4G). This contrast quantifies, at scale, the qualitative picture that emerges from individual structures: amyloid assemblies are stabilized by compact cooperative regions embedded within a matrix of constrained, locally frustrated elements.

Finally, to obtain an interpretable, low-dimensional view that connects residue composition, energetic role, and variability, we constructed a two-dimensional energy– variability landscape by classifying residues according to the global medians of their mean ΔG and variance. This yields four quadrants: stable-invariant, stable-variable, neutral/destabilizing-invariant, and neutral/destabilizing-variable, that partition residues by energetic contribution and plasticity (Fig. 4H). Quadrant-specific amino-acid enrichments recapitulate intuitive biochemistry: hydrophobic residues concentrate within the stable quadrants, whereas polar and charged residues are overrepresented in destabilizing or variable regimes (Fig. 4I). Aggregating the per-protein fractions of residues in each quadrant (together with disorder content) produces compact descriptors that separate proteins and functional classes by coarse-grained energetic architecture. This complements traditional sequence- or topology-based classifications by emphasizing the thermodynamic roles residues play within the fibril context. Across all analyses, the same theme emerges: standardized, structure-embedded energetics enable principled alignment with physicochemical scales, quantification of protein-level heterogeneity, discrimination between cooperative stabilization and local frustration by characteristic length scales, and construction of interpretable energetic descriptors for classification and prediction. This global synthesis illustrates the central motivation for Amyloid Explorer—turning a rapidly growing structural archive into a platform for mechanism, taxonomy, and design.

## Discussion

Amyloid Explorer establishes a comprehensive and rigorously annotated resource to date for the comparative analysis of amyloid fibril structures. By combining large-scale structural curation with systematic residue-level and cooperative thermodynamic profiles, structural quality metrics, and provenance metadata, the platform provides a foundation for reproducible, scalable interrogation of amyloid polymorphism. The inclusion of real-time visualization tools and programmatic access ensures that both exploratory and automated workflows can access and interrogate the dataset efficiently.

The resource builds on previous frameworks by combining structural curation with energetic interpretation, allowing users not only to visualize fibril architecture, but also to quantify stabilization patterns, identify aggregation-prone regions, and compare fibril variants under different experimental conditions. These features enable mechanistic dissection of fibril formation, mutation effects, and structural class differences, and are applicable to both hypothesis-driven and large-scale statistical analyses^5,16-19,21,23,24^.

The global patterns uncovered here argue that, despite extensive sequence diversity and experimental variation, amyloid fibrils are shaped by a constrained thermodynamic framework. Cooperative APRs provide compact stabilizing cores; context-dependent rearrangements of the same cores yield divergent topologies (framework polymorphism); and short, spatially organized frustrated segments delimit flexibility at boundaries and interfaces. Residue-level anchoring minima provide energetic fingerprints for strains and align with mutational hotspots in disease, offering a mechanistic explanation for how small sequence changes can rewire stabilization patterns and, consequently, strain identity^24,25^. The inverse relationship between sequence complexity and polymorphic breadth further suggests that low-complexity sequences afford greater packing plasticity, thereby facilitating alternative fold solutions.

Beyond classification, this constrained thermodynamic framework offers practical applications. It supports prediction of aggregation-prone segments and polymorph behavior, guides interpretation of disease-linked mutations, and can inform the design of aggregation inhibitors or stabilizing modulators^20,45-48^. The accumulation of fibril structures now mirrors the expansion of globular protein structures in the 1980s, when systematic comparison enabled the creation of fold classification systems such as SCOP^49^ and CATH^50^. Those resources transformed structural biology into a predictive science by distilling general architectural rules from a growing archive. Amyloid Explorer provides a similar inflection point for the amyloid field: by unifying hundreds of fibril models under a single annotated framework, it establishes the foundation for a principled taxonomy of amyloid folds and their underlying energetic logic. As more fibril structures become available, particularly from functional amyloids or under perturbed cellular conditions, Amyloid Explorer provides a ready infrastructure for comparative annotation and hypothesis testing at scale.

We anticipate that these extensions will progressively transform the resource from a static atlas into a dynamic engine beyond serving structural biologists, to clinicians, medicinal chemists and data scientists for mechanistic discovery, classification, and therapeutic modulation.

## Materials and Methods

### Retrieval of amyloid fibril structures

Amyloid Explorer integrates structural data from more than six hundred experimentally determined amyloid fibril structures across more than 50 unique protein sequences. Structures were retrieved from the Protein Data Bank (PDB)^12^, Electron Microscopy Data Bank (EMDB)^13^, and Biological Magnetic Resonance Data Bank (BMRB)^14^ using targeted queries for amyloid assemblies. This dataset reflects the full scope of currently available high-resolution amyloid models. For consistency, all models were transformed to contain a standard number of stacked monomers (10) and uniformly renumbered to align with database parsing and comparative analysis tools. Each protofilament chain was renamed in a systematic fashion and stack directions standardized.

### Data and web interface navigation

Amyloid Explorer’s construction followed a multi-step process involving the retrieval and annotation of fibril structures, integration of structural, thermodynamic, and metadata properties, development of visualization and analysis tools, thorough data validation, and the deployment of a scalable, cloud-hosted web interface. The entire platform is hosted using a remote-first architecture on the Netlify cloud service, ensuring high availability and responsive performance. All data are stored in a structured format on a dedicated server environment and are freely available, requiring no registration for access. Importantly, the legacy STAMP-dB URL (https://stampdb.switchlab.org) remains active and redirect to the updated entries within Amyloid Explorer, preserving continuity.

### Structural quality control

To ensure data reliability and consistency, every fibril model structure underwent rigorous quality control analysis. Resolution was recorded directly from PDB or EMDB metadata, and models with poor resolution were flagged. We defined an empirical partitioning boundary based on a linear relationship between structural resolution and van der Waals (VDW) clashes using the equation:

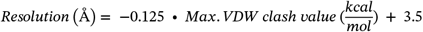

This threshold reveals models with both poor resolution and high steric clashes that can affect downstream analysis. The resolution values used are the original reported in the source publications, determined by the Fourier Shell Correlation (FSC) 0.143 criterion^38^, which estimates the resolution of the reconstructed cryo-EM maps. Van der Waals clashes were identified using FoldX^39^. Backbone dihedral statistics (φ/ψ angles) and Ramachandran outliers were determined using in-house scripts and PyRosetta^37^. Heterotypic components of the structures (e.g., detergents, polyanionic co-factors) were removed as they could not be used to calculate energetics.

### Thermodynamic annotation

Thermodynamic annotation was carried out using two complementary approaches. First, per-residue ΔG values were computed using FoldX^39^, based on the residue’s contributions to protofilament integrity and cross-β interactions. The structures were energetically optimized using the FoldX *RepairPDB* function and individual residue energetics were calculated using the *SequenceDetail* function. The energies of each residue were then averaged at the level of individual protofilaments and across each protofilament stack, excluding the residues corresponding to the exposed monomers on each side of the stack, to account for the lack of additional contacts at the fibril tips.

For the window mode, we adapted from a recent protocol. Briefly, applying a sliding window mode (5-residues per window), we averaged the values over a sliding window of size 5 and linearly rescaled the values to a mean of 0 and standard deviation of 1. This method allows identification of cooperative aggregation-prone regions (APRs), using a previously established methodology^17^. Both views are provided through an interactive toggle interface on the database.

### Global sequence–energetics analyses

Per-residue energetic records—comprising protein and structure identifiers, residue type and index, and ΔG terms including raw individual residue energies (Average_Energy_Stack_1) and window energies (Avg_SW_Energy_Stack_1), were aggregated across all standardized structures. Amino-acid physicochemical property scales were obtained from AAindex^43^.

For scale associations, the mean energetic contribution of each aminoacid type was correlated with the corresponding scale values using both Pearson and Spearman metrics, with Benjamini–Hochberg correction applied across scales. The energy-diversity score for each protein was defined as the mean standard deviation of per-residue energies across all structures available for that protein; visual encodings indicate the number of structures contributing to each estimate.

For segment analyses, sign-coherent runs of negative (stabilizing) and positive (destabilizing) ΔG were enumerated after mild smoothing to reduce single-site noise, and length distributions were compared using non-parametric tests where appropriate. For the energy–variability landscape, residues were assigned to quadrants defined by the global medians of mean ΔG and variance; amino-acid enrichments were computed for each quadrant. Protein-level descriptors comprising quadrant fractions (plus disorder fraction) were embedded into a low-dimensional space to visualize relationships among proteins and functional classes.

All analyses were performed in R using the tidyverse and ggplot2 libraries for visualization, with harmonized themes and aspect ratios to facilitate cross-panel comparison.

## Acknowledgements

The Switch Laboratory was supported by the Flanders Institute for Biotechnology (VIB, grant no. C0401 to FR and JS), the Fund for Scientific Research Flanders (FWO, project grant G0A6724N to JS), and the Stichting Alzheimer Onderzoek /Fondation Recherche Alzheimer (project grant SAO-FRA 2023/0005 to JS). NL was supported by a Thomas O. Hicks Endowed Scholarship. Computational resources were provided by the BioHPC cluster supported by the Lyda Hill Department of Bioinformatics at UTSW.

## Author contributions

All authors contributed to data collection. VK and NL performed the thermodynamic analysis of amyloid fibril structures. NL, KK, and LG annotated the dataset. NL, JS, and FR performed meta-analysis. NL, JS, and FR acquired funding and supervised the project. NL, JS, and FR prepared the original draft and all authors contributed to the review and editing of the final version of this manuscript.

## Data Availability statement

Amyloid Explorer is publicly available online via its new dedicated website (https://amyloid-explorer.switchlab.org), as well as through the original STAMP-dB address (https://stamp.switchlab.org). The database and its contents are freely accessible without registration or usage restrictions.

## References

1 Buxbaum, J. N. et al. Amyloid nomenclature 2024: update, novel proteins, and recommendations by the International Society of Amyloidosis (ISA) Nomenclature Committee. Amyloid 31, 249–256 (2024).

2 Chiti, F. & Dobson, C. M. Protein Misfolding, Amyloid Formation, and Human Disease: A Summary of Progress Over the Last Decade. Annu Rev Biochem 86, 27–68 (2017).

3 Louros, N., Schymkowitz, J. & Rousseau, F. Mechanisms and pathology of protein misfolding and aggregation. Nat Rev Mol Cell Biol 24, 912–933 (2023).

4 Otzen, D. & Riek, R. Functional Amyloids. Cold Spring Harb Perspect Biol 11 (2019).

5 Otzen, D. E. et al. Interactions between pathological and functional amyloid: A match made in Heaven or Hell? Mol Aspects Med 103, 101351 (2025).

6 Ledvina, H. E. et al. Functional amyloid proteins confer defence against predatory bacteria. Nature (2025).

7 Fitzpatrick, A. W. P. et al. Cryo-EM structures of tau filaments from Alzheimer’s disease. Nature 547, 185–190 (2017).

8 Scheres, S. H. W., Ryskeldi-Falcon, B. & Goedert, M. Molecular pathology of neurodegenerative diseases by cryo-EM of amyloids. Nature 621, 701–710 (2023).

9 Nguyen, B. A. et al. Structural polymorphism of amyloid fibrils in ATTR amyloidosis revealed by cryo-electron microscopy. Nature Communications 15, 581 (2024).

10 Radamaker, L. et al. Cryo-EM reveals structural breaks in a patient-derived amyloid fibril from systemic AL amyloidosis. Nat Commun 12, 875 (2021).

11 Shi, Y. et al. Structure-based classification of tauopathies. Nature 598, 359–363 (2021).

12 Burley, Stephen K. et al. Updated resources for exploring experimentally-determined PDB structures and Computed Structure Models at the RCSB Protein Data Bank. Nucleic Acids Research 53, D564–D574 (2025).

13 The ww, P. D. B. C. EMDB—the Electron Microscopy Data Bank. Nucleic Acids Research 52, D456–D465 (2024).

14 Hoch, J. C. et al. Biological Magnetic Resonance Data Bank. Nucleic Acids Research 51, D368–D376 (2023).

15 Gallardo, R., Ranson, N. A. & Radford, S. E. Amyloid structures: much more than just a cross-β fold. Curr Opin Struct Biol 60, 7–16 (2020).

16 Louros, N., Schymkowitz, J. & Rousseau, F. Heterotypic amyloid interactions: Clues to polymorphic bias and selective cellular vulnerability? Curr Opin Struct Biol 72, 176–186 (2022).

17 van der Kant, R., Louros, N., Schymkowitz, J. & Rousseau, F. Thermodynamic analysis of amyloid fibril structures reveals a common framework for stability in amyloid polymorphs. Structure 30, 1178–1189.e1173 (2022).

18 Louros, N., Schymkowitz, J. & Rousseau, F. Toward a thermodynamic taxonomy of amyloid fibrils. Structure 33, 1628–1630 (2025).

19 Mullapudi, V. et al. Network of hotspot interactions cluster tau amyloid folds. Nature Communications 14, 895 (2023).

20 Louros, N. et al. Mapping the sequence specificity of heterotypic amyloid interactions enables the identification of aggregation modifiers. Nature Communications 13, 1351 (2022).

21 Duran-Romaña, R., Schymkowitz, J., Rousseau, F. & Louros, N. Energetic profiling reveals thermodynamic principles underlying amyloid fibril maturation. bioRxiv, 2025.2005.2014.653959 (2025).

22 Louros, N. et al. Local structural preferences in shaping tau amyloid polymorphism. Nat Commun 15, 1028 (2024).

23 Arutyunyan, A., Seuma, M., Faure, A. J., Bolognesi, B. & Lehner, B. Massively parallel genetic perturbation suggests the energetic structure of an amyloid-β transition state. Science Advances 11, eadv1422

24 Connor, J. P., Radford, S. E. & Brockwell, D. J. Structural and thermodynamic classification of amyloid polymorphs. Structure (2025).

25 Louros, N., Schymkowitz, J. & Rousseau, F. Mechanisms and pathology of protein misfolding and aggregation. Nature Reviews Molecular Cell Biology 24, 912–933 (2023).

26 Sawaya, M. R., Hughes, M. P., Rodriguez, J. A., Riek, R. & Eisenberg, D. S. The expanding amyloid family: Structure, stability, function, and pathogenesis. Cell 184, 4857–4873 (2021).

27 Louros, N. et al. WALTZ-DB 2.0: an updated database containing structural information of experimentally determined amyloid-forming peptides. Nucleic Acids Research 48, D389–D393 (2020).

28 Varadi, M., De Baets, G., Vranken, W. F., Tompa, P. & Pancsa, R. AmyPro: a database of proteins with validated amyloidogenic regions. Nucleic Acids Res 46, D387–d392 (2018).

29 Rawat, P. et al. CPAD 2.0: a repository of curated experimental data on aggregating proteins and peptides. Amyloid 27, 128–133 (2020).

30 Louros, N., van der Kant, R., Schymkowitz, J. & Rousseau, F. StAmP-DB: a platform for structures of polymorphic amyloid fibril cores. Bioinformatics 38, 2636–2638 (2022).

31 Yanagisawa, H. et al. Cryo-EM of wild-type and mutant PMEL amyloid cores reveals structural mechanism of pigment dispersion syndrome. Nat Commun 16, 5411 (2025).

32 Flores, M. D. et al. Structure of a reversible amyloid fibril formed by the CPEB3 prion-like domain reveals a core sequence involved in translational regulation. bioRxiv, 2022.2012.2007.519389 (2022).

33 Hervas, R. et al. Cryo-EM structure of a neuronal functional amyloid implicated in memory persistence in Drosophila. Science 367, 1230–1234 (2020).

34 Bücker, R. et al. The Cryo-EM structures of two amphibian antimicrobial cross-β amyloid fibrils. Nature Communications 13, 4356 (2022).

35 Nagy-Smith, K., Moore, E., Schneider, J. & Tycko, R. Molecular structure of monomorphic peptide fibrils within a kinetically trapped hydrogel network. Proceedings of the National Academy of Sciences 112, 9816–9821 (2015).

36 Rose, A. S. & Hildebrand, P. W. NGL Viewer: a web application for molecular visualization. Nucleic Acids Research 43, W576–W579 (2015).

37 Chaudhury, S., Lyskov, S. & Gray, J. J. PyRosetta: a script-based interface for implementing molecular modeling algorithms using Rosetta. Bioinformatics 26, 689–691 (2010).

38 Rosenthal, P. B. & Henderson, R. Optimal determination of particle orientation, absolute hand, and contrast loss in single-particle electron cryomicroscopy. J Mol Biol 333, 721–745 (2003).

39 Schymkowitz, J. et al. The FoldX web server: an online force field. Nucleic Acids Res 33, W382–388 (2005).

40 Ramachandran, G. & Udgaonkar, J. B. Difference in Fibril Core Stability between Two Tau Four-Repeat Domain Proteins: A Hydrogen–Deuterium Exchange Coupled to Mass Spectrometry Study. Biochemistry 52, 8787–8789 (2013).

41 Illes-Toth, E., Rempel, D. L. & Gross, M. L. Pulsed Hydrogen– Deuterium Exchange Illuminates the Aggregation Kinetics of α-Synuclein, the Causative Agent for Parkinson’s Disease. ACS Chemical Neuroscience 9, 1469–1476 (2018).

42 Wälti, M. A., Orts, J. & Riek, R. Quenched hydrogen-deuterium exchange NMR of a disease-relevant Aβ(1-42) amyloid polymorph. PLOS ONE 12, e0172862 (2017).

43 Kawashima, S. et al. AAindex: amino acid index database, progress report 2008. Nucleic Acids Res 36, D202–205 (2008).

44 Conchillo-Solé, O. et al. AGGRESCAN: a server for the prediction and evaluation of “hot spots” of aggregation in polypeptides. BMC Bioinformatics 8, 65 (2007).

45 Konstantoulea, K. et al. Heterotypic Amyloid beta interactions facilitate amyloid assembly and modify amyloid structure. EMBO J 41, e108591 (2022).

46 Wagner, J. et al. Medin co-aggregates with vascular amyloid-beta in Alzheimer’s disease. Nature 612, 123–131 (2022).

47 Murray, K. A. et al. Identifying amyloid-related diseases by mapping mutations in low-complexity protein domains to pathologies. Nature Structural & Molecular Biology 29, 529–536 (2022).

48 Sønderby, T. V. et al. Sequence-targeted Peptides Divert Functional Bacterial Amyloid Towards Destabilized Aggregates and Reduce Biofilm Formation. J Mol Biol 435, 168039 (2023).

49 Fox, N. K., Brenner, S. E. & Chandonia, J.-M. SCOPe: Structural Classification of Proteins—extended, integrating SCOP and ASTRAL data and classification of new structures. Nucleic Acids Research 42, D304–D309 (2014).

50 Knudsen, M. & Wiuf, C. The CATH database. Hum Genomics 4, 207–212 (2010).

